# Inhibition of Tau seeding by targeting Tau nucleation core within neurons with a single domain antibody fragment

**DOI:** 10.1101/2021.03.23.436266

**Authors:** Clément Danis, Elian Dupré, Orgeta Zejneli, Raphaëlle Caillierez, Alexis Arrial, Séverine Bégard, Anne Loyens, Justine Mortelecque, François-Xavier Cantrelle, Xavier Hanoulle, Jean-Christophe Rain, Morvane Colin, Luc Buée, Isabelle Landrieu

## Abstract

Tau proteins aggregate into filaments in brain cells in Alzheimer’s disease and related disorders referred to as tauopathies. Here, we used fragments of camelid heavy-chain-only antibodies (VHHs or single domain antibody fragments) targeting Tau as immuno-modulators of its pathologic seeding. A VHH issued from the screen against Tau of a synthetic phage-display library of humanized VHHs was selected for its capacity to bind Tau microtubule-binding domain, composing the core of Tau fibrils. This lead VHH was optimized to improve its biochemical properties and to act in the intracellular compartment, resulting in VHH Z70. VHH Z70 was more efficient than the lead to inhibit *in vitro* Tau aggregation in heparin-induced assays. Expression of VHH Z70 in a cellular model of Tau seeding also decreased the fluorescence-reported aggregation. Finally, intracellular expression of VHH Z70 in the brain of an established tauopathy mouse seeding model demonstrated its capacity to mitigate accumulation of pathological Tau. VHH Z70, by targeting Tau inside brain neurons, where most of the pathological Tau resides, provides a new tool to explore the optimal strategies of immunotherapy in tauopathies.

## Introduction

Aggregation of the intrinsically disordered neuronal Tau protein to form fibrillar amyloid structures is related to neurodegenerative disorders called tauopathies, including the most prevalent, Alzheimer’s disease (AD). AD is characterized by both extracellular amyloid deposits made of Aß (amyloid) peptides and intraneuronal neurofibrillary tangles (NFTs) formed by Tau protein aggregates^1^. Intervention strategies based on the amyloid cascade hypothesis had, up to date, limited success despite their being the primary target of clinical assays^2^. In AD, the severity of cognitive decline is better correlated with the evolution of NFTs than amyloid deposits^3–5^. In other tauopathies, no amyloid deposition is observed. This emphasizes the need to pursue other biological hypotheses, including the Tau pathway, in search for disease-mitigating treatments for tauopathies.

In the pathological context, Tau is the principal component of paired helical filaments (PHFs) and straight filaments^6,7^, which form the intracellular fibrillar deposits leading to the NFTs and ultimately to neurofibrillary degeneration. In addition, it has been proposed that extracellular pathological Tau are taken up in cells, leading to intracellular Tau seeding of the aggregation process^8–10^. The 441-amino acid residues longest Tau isoform can be divided into 4 domains comprising the N-terminal domain (N1-N2), the proline-rich domain (P1-P2), the microtubule-binding domain (MTBD) constituted itself of 4 partially repeated regions, R1 to R4, and the C-terminal domain (**Fig. 1b** scheme). Two homologous hexapeptides named PHF6* (_275_VQIINK_280_) and PHF6 (_306_VQIVYK_311_) located respectively at the beginning of R2 and R3 repeat regions (**Fig. 1b** scheme) of Tau MTBD are nuclei of Tau aggregation^11^. PHF6* and PHF6 peptides spontaneously aggregate in solution contrary to the full-length Tau that is a highly soluble protein. Their atomic structures reveal the capacity of these segments to form interdigitated steric-zipper interfaces that seed Tau aggregation^12,13^. The structures of Tau fibers isolated from patient brains affected by various tauopathies: AD^14^, corticobasal degeneration^15^, Pick’s disease^16^ and chronic traumatic encephalopathy^17^ were resolved by cryo-electron microscopy. The common core of these fibrillary structures is composed of the subdomains R3 including the PHF6, R4 and a part of the C-terminal domain (V306-F378) that mainly form a β-sheet structure^14^.

**Fig 1.**
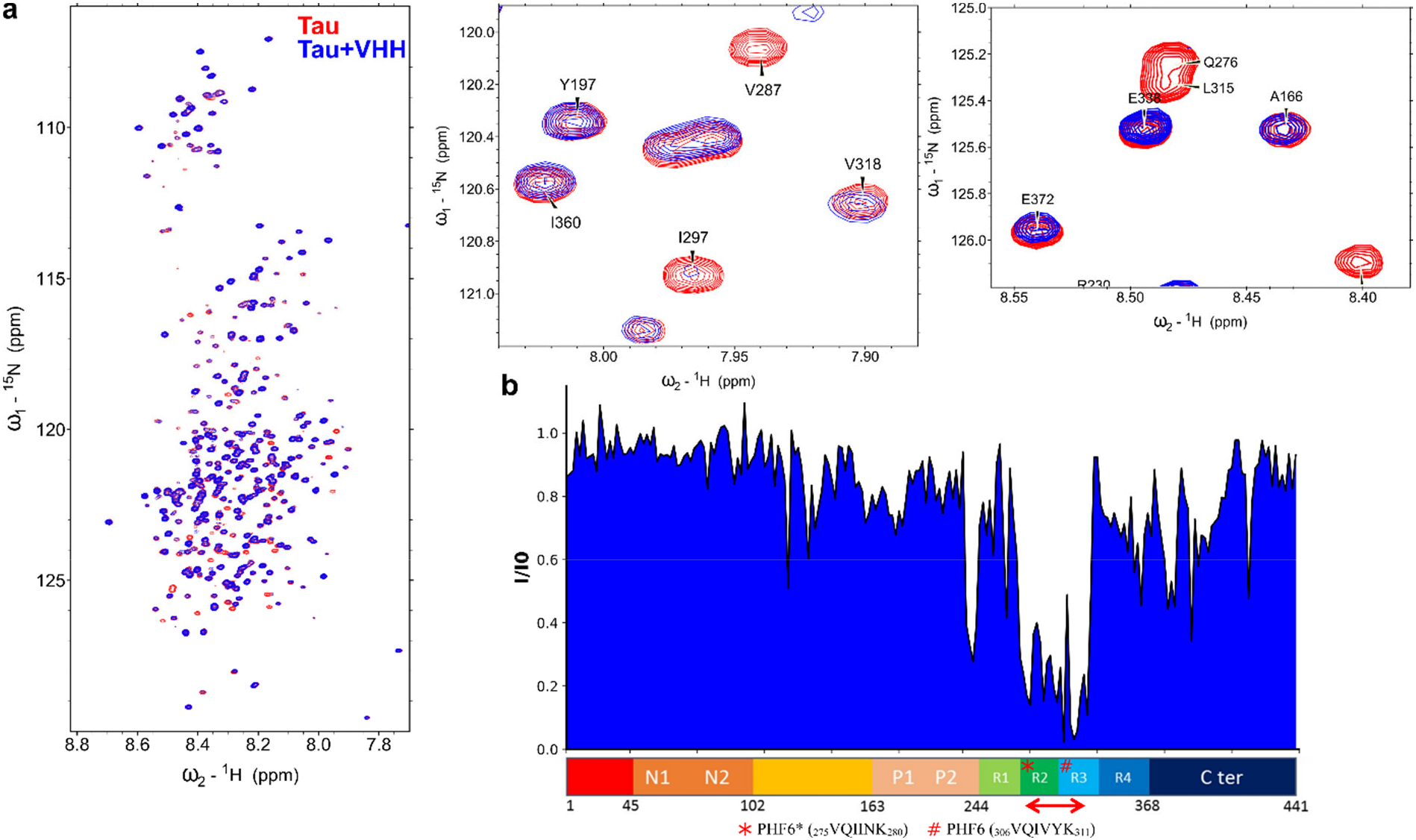
VHH E4-1 binds to the MTBD of Tau **a** Overlay of two-dimensional ^1^H, ^15^N HSQC spectra of Tau (in red) with Tau mixed with non-labelled VHH E4-1 (superimposed in blue) (n=1). In the spectrum of Tau in the presence of VHH E4-1, multiple resonances are broadened beyond detection compared to the Tau control spectrum. Spectra enlargements evidence broadened resonances corresponding to residues implicated in the interaction. Normalized NMR intensities (I/I0) along the Tau sequence with (I0) and (I) corresponding to the resonance intensity when Tau is free in solution or mixed with equimolar quantity of VHH E4-1 (I), respectively. The normalized intensity ratios (I/I0) plot allowed the identification of the Tau MTBD domain as the target of VHH E4-1 interaction. Overlapping resonances are not considered (*x-axis* is not scaled). A red double-arrow indicates the region containing the corresponding major broadened resonances, which was mapped to the R2-R3 repeats in the MTBD.

The compelling results of immunotherapies directed against Tau in several transgenic (Tg) mouse models of tauopathies, in decreasing Tau accumulation and in some case ensuring recovery of cognitive or motor functions^18–25^, have motivated several pharmaceutical companies to launch clinical trials of active and passive immunization, the latter with various anti-Tau monoclonal antibodies^26,27^ (Immunoglobulin G, IgG). However, advances in the field still require deciphering key aspects of efficient anti-Tau immunotherapies and to develop their full potential to target tauopathies. The road to anti-Tau immunotherapies opened based on the evidence of Tau seeding capacity and propagation between neurons^28–31^. Yet, the majority of pathological Tau remains intracellular in the cytoplasm, where it is not the primary target of anti-Tau conventional immunotherapies using IgG. In addition, the extracellular Tau could, at least partly, remains unattainable to antibodies, as would be the case for Tau in extracellular vesicles^32^ or nanotubes^33^. Finally, the propagation pattern related to extracellular Tau, clearly defined in AD, could also be less relevant in pure and more acute tauopathies, for which the time frame restrains the spreading^30^. In these latter cases, the rational to target extracellular Tau is weaker. However, the multiple trials in mouse models with IgGs repeatedly showed their efficacy, despite most approaches limited to targeting the extracellular Tau^18–24^. Nevertheless, there is little documentation on their capacity to act in the intraneuronal compartment, following uptake, a mode of action that should be accounted for at least some of them^34^. One study with systematic comparison of intraneuronal *versus* extraneuronal-acting equivalent anti-Tau scFvs (single chain fragment of the variable region of IgG) showed a significant improved effect of the intraneuronal modified scFvs against the tauopathy, in two tauopathy mouse models^35^. Similarly, comparison of IgG 6B2 and intraneuronal modified scFv 4E6, which share the same Tau-binding loops, has shown that only scFv 4E6 is effective at mitigating the tauopathy in cellular and *in vivo drosophila* models^36^. Intraneuronal chimeric scFvs are another successful example, designed to target Tau to the proteasome or lysozomal pathway in human Tau Tg mice^37^.

Exploring the capacity to use immunotherapies targeting the intraneuronal Tau has thus become an important challenge to further explore. With that in mind, we have chosen VHHs (Variable Heavy-chain of the Heavy-chain-only antibody), commonly called nanobodies, to target new and original epitopes of Tau, and to optimize their intracellular activity. VHHs consist in an unique heavy-chain that corresponds to the variable heavy-chain from *Camelidae* immunoglobulin G^38^. The interest of using the VHHs instead of the classical antibodies stand in their easy generation, from a synthetic library, involving no animal handling, their selection using phage-display, their production in periplasm of bacteria, as well as the multiple possibilities offered by modification using protein engineering^39^. They can be modified to penetrate into the cytoplasm of cells, or expressed inside the cells, and bind specifically to their target epitope^40,41^. Additionally, VHHs showed their potential as diagnostic tools to target NFTs with an affinity and specificity very close to antibodies already used for detecting these pathologic features by immunochemistry, opening the way for new probes in *in vivo* imaging experiments^42^. We here demonstrated that it is also possible to consider VHHs as therapeutic tools in tauopathies.

We generated, optimized and characterized a VHH targeted against Tau MTBD, obtained by screening from a humanized naive synthetic library. The optimized version of this lead VHH, named Z70, inhibits Tau aggregation *in vitro* and in HEK293 seeding-reporting cellular model. Lentiviral vector (LV) expression of VHH Z70 inside brain cells in the hippocampus area of a tauopathy mouse model, blocked the seeding of the pathology induced by human AD seeds. Z70 antibody fragment thus provides a new tool to define the best strategies in Tau immunotherapies and to open the way for future gene therapy.

## Results

### Identification of a synthetic VHH directed against Tau microtubule-binding domain

To generate original VHHs targeting Tau, with the potential to block its aggregation, 20 clones resulting from the screen of a synthetic phage-display library of humanized llama single-domain antibody^43^ against recombinant Tau protein (Tau 2N4R longest isoform) were initially selected for further analysis. Definition of the epitope recognized by each of these VHHs was a first step in assessing their properties. Resonance perturbation mapping in ^1^H, ^15^N HSQC spectra of ^15^N-Tau, obtained by nuclear magnetic resonance (NMR) spectroscopy, allowed to define the various binding sites of the VHHs along Tau sequence. Interaction results in spectral perturbations by modifying the chemical environment and/or the local dynamics near the binding site, which can thus be defined by comparison of the spectra of Tau alone in solution or in the presence of a VHH. Each resonance in Tau spectra can be linked to a specific amino acid residue in Tau sequence^44,45^, allowing to map the binding site based on the observed differences of chemical shift value or intensity for each resonance in these bound/free conditions. Interestingly, one VHH, named VHH E4-1, affected resonances in Tau spectrum corresponding to residues in the MTBD (**Fig. 1**). The epitope mapping was refined using a Tau fragment that corresponded to the isolated MTBD. The smaller size of this Tau fragment (124 amino acid residues instead of 441) resulted in fewer resonances and less resonance overlap in the corresponding Tau[245-368] ^1^H, ^15^N spectrum, facilitating the identification of the binding site (**Supplementary Figs. 1-2**). The affected resonances corresponded to amino acid residues located in a stretch expanding from residue V275 to K317 (**Supplementary Fig. 2b**). VHH E4-1 thus bound within the R2-R3 repeats of the MTBD.

### Optimization of lead VHH E4-1 into variant VHH Z70

An important property of the VHH that we wished to develop is its capacity to be expressed and to recognize its target in the cytoplasmic environment, inside the cells. However, some VHHs might be ineffective in binding once expressed in a cell, due to improper folding and/or poor stability. Indeed, VHH E4-1 proved to be a poor binder of Tau in yeast two-hybrid assays that require the interaction to take place in the yeast nucleus, providing an evaluation of VHH E4-1 intracellular binding capacity^46,47^ (**Fig. 2a**).VHH E4-1 was thus next submitted to a round of optimization, using yeast two-hybrid system, to maximize its capacity to recognize its target when expressed in a cellular environment. First, we built a cDNA mutant library by random mutagenesis, targeting the whole sequence of VHH E4-1 to produce a variety of VHH preys (C-terminal Gal4-activation domain fusion) against the Tau bait (N-terminal LexA fusion). The library was transformed in yeast and screen by cell-to-cell mating to get positive colonies under the pressure of selection conditions corresponding to undetected VHH E4-1-Tau interaction (growth medium lacking amino acids Leu, Trp and His, **Fig. 2a**). An optimized variant, named VHH Z70, was selected, resulting from 4 mutations G12V, P16S, T81M and W114G located in the framework domains or FR, outside the recognition loops or CDR (**Fig. 2a-b**), suggesting that the epitope recognized by this mutant is unaltered. Conservation of the epitope was confirmed by NMR resonance perturbation mapping, using labelled Tau and MTBD in the same manner as for the lead VHH E4-1 (**Supplementary Figs. 3-4**). Interaction of VHH E4-1 and VHH Z70 with Tau were further characterized using surface plasmon resonance spectroscopy (SPR) with biotinylated-Tau immobilized at the surface of a streptavidin-functionalized chip. The assay provided the kinetic parameters of the interaction, characterized by dissociation constants Kd of 345 nM for VHH E4-1 (**Fig. 2c**) and Kd of 147 nM for variant VHH Z70 (**Fig. 2d**). VHH Z70, optimized for intracellular activity, had a better affinity for its target than VHH E4-1, the major optimization concerning the association constant (*k*_on_) (**Supplementary Fig. 5**). SPR was additionally performed with VHH Z70 immobilized on the chips. A Tau peptide [273-318] corresponding to the NMR-identified VHH binding site, fused to a SUMO domain to improve its solubility, was injected into the flux. VHH Z70 interacted with the fused peptide with a Kd of 85 nM (**Fig. 2e, Supplementary Fig. 5**), confirming that the region V275-K317 in Tau sequence was self-sufficient for VHH-Z70 binding.

**Fig 2.**
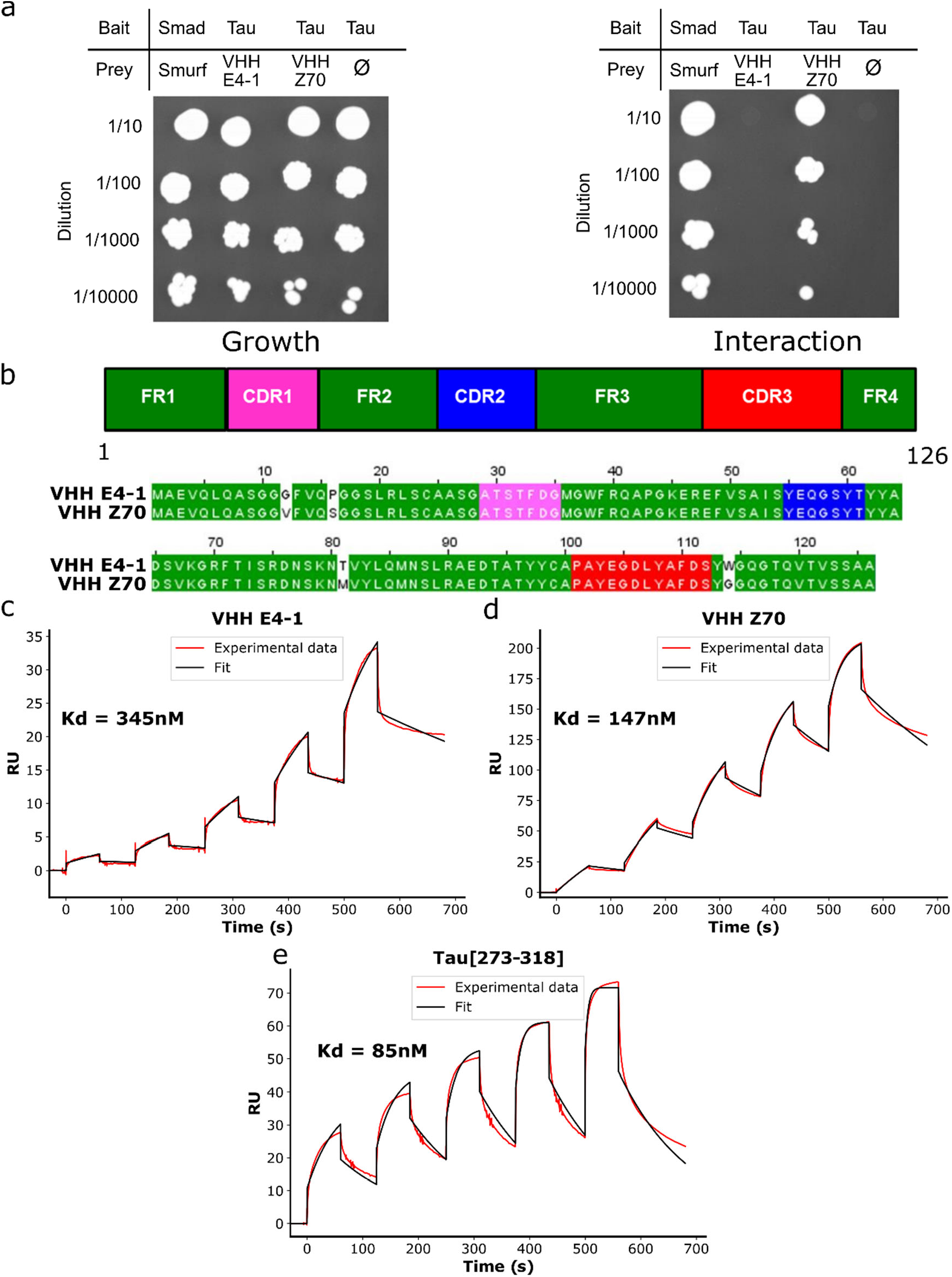
VHH Z70 is optimized for intracellular binding and has a better affinity for Tau than VHH E4-1 **a**: Results from yeast two-hybrid. A growth test on non-selective medium (*left panel*, lacking only leucine and tryptophane) or on selective medium (*right panel*, lacking leucine, tryptophane and histidine) was performed with dilution (*top to bottom*) of the diploid yeast culture expressing both bait and prey constructs. Positive and negative controls of interaction consist respectively in Smad/Smurf interaction^71^and Tau alone (empty vector). VHH E4-1 did not interact with Tau in yeast (no growth on selective medium) whereas VHH Z70 did. **b**: Domain organization of the VHHs (CDR are for complementarity-determining regions and FR for framework domains) and sequence alignment between VHH E4-1 and VHH Z70 showing 4 mutations in the FR: G12V, P16S, T81M and W114G. **c, d** Sensorgrams (reference subtracted data) of single cycle kinetics analysis performed on immobilized biotinylated Tau, with five injections of VHH E4-1 (**c**) or VHH Z70 (**d**) at 0.125 µM, 0.25 µM, 0.5 µM, 1 µM, and 2 µM (n=1). Dissociation equilibrium constant Kd were calculated from the ratio of off-rate and on-rate kinetic constants k_off_/k_on_ (**Supplementary Fig. 5**). **e** Sensorgram (reference subtracted data) of single-cycle kinetics analysis performed on immobilized VHH Z70 on a CM5 chip, with five injections of peptide Tau[273-318] fused at its N-terminus with the SUMO protein (n=1). Tau peptide sequence and, k_on_, k_off_ and Kd are included in the table in **Supplementary Fig. 5**. Black lines correspond to the fitted curves, red lines to the measurements.

### Identification of the minimal Tau epitope recognized by VHH Z70

The binding site identified by NMR for both lead VHH E4-1 and optimized VHH Z70 was larger than expected for an epitope, about 45 contiguous amino-acid residues showing strong reduction of their resonance intensities upon binding. To determine the minimal epitope that VHH Z70 can recognize, an epitope mapping was performed using yeast two-hybrid (267.10^3^ tested interactions) with VHH Z70 as bait (LexA-VHH fusion) against a library of Tau fragments as preys (GAL4_activation domain-Tau_fragments). 90 positive clones were selected for their growth in the selection conditions, evidencing binding of VHH Z70 to Tau fragments of various length, from a small-scale cell-to-cell mating screen.

Comparison of the Tau prey fragment sequences corresponding to these 90 interactions identified peptide _305_SVQIVYKPV_313_ as the minimal common recognition motif of Tau that VHH Z70 can bind (**Supplementary Fig. 6**). The sequence is localized in the R3 repeat of the MTBD domain and contains the PHF6 peptide _306_VQIVYK_311_. We next used Tau2N3R isoform, which lacks the R2 repeat and so does not contain the PHF6* peptide, to confirm that the R3 repeat, containing the PHF6 peptide, was sufficient for the interaction. As observed in the resonance intensity profile, the interaction of VHH Z70 with Tau2N3R is maintained, and the most affected resonances in the Tau spectrum corresponded to the PHF6 residues in the R3 repeats (**Supplementary Fig. 7**), confirming that PHF6* is not necessary for VHH Z70 binding to Tau.

### Inhibition of *in vitro* Tau aggregation

VHH E4-1 and VHH Z70 recognizing Tau peptide PHF6, known to nucleate the aggregation and to form the core of Tau fibers, were assayed for their capacity to interfere with Tau *in vitro* aggregation. The assays were carried out with Tau recombinant protein in the presence of heparin, using thioflavin T as a dye whose fluorescence is increased in presence of aggregates **(Fig. 3**). Negative and positive controls consisted in Tau without or with heparin, respectively. An additional control was performed in the presence of VHH F8-2, a VHH issued from the initial phage-library screen, which targets Tau C-terminal domain^48^. At 10µM of Tau, the observed amount of aggregates was maximal (defined as 100%) for the positive control after 8 h of incubation at 37°C, while no fluorescence change was detected for the negative control (**Fig. 3a-c**). At equimolar concentration of Tau:VHH F8-2, the fluorescence signal reached 91,2 % (± 3.8%), showing that VHH F8-2 did not affect the aggregation of Tau (**Fig. 3a**). In contrast, at a molar ratio of 1:0.25 Tau:VHH E4-1, the maximal fluorescence signal reached 86,9 % (± 2.4%). Additionally, about 3.8h were needed to gain 50% of maximal signal, compare to 2.5h for the positive control, showing a slower aggregation kinetic in the presence of VHH E4-1 (**Fig. 3b**). At a 1:1 Tau:VHH E4-1 molar ratio, the fluorescence signal only reached 58.3 % (± 3.9%) and more than 12.8h were necessary to gain 50% of maximal signal (**Fig. 3b**). VHH Z70 had an even stronger inhibition effect on the aggregation of Tau than the lead VHH E4-1. At a 1:1 Tau:VHH Z70 molar ratio, the maximal fluorescence signal barely reached above the negative control level, at 4.1 % (± 0.1%) (**Fig. 3c**). The link between the thioflavin T fluorescence measurements in our assays and the formation of Tau aggregates at the end-point of each aggregation assay was confirmed by transmission electron microscopy imaging that allowed direct visualization of typical Tau fibers, whether present (**Fig. 3d-h**). Large amounts of fibrils were observed for Tau in the presence of heparin only (**Fig. 3e**) or in the additional presence of VHH F8-2 (**Fig. 3f**), but shorter filaments with VHH E4-1 (**Fig. 3g**) and practically none with VHH Z70 (**Fig. 3h**). In conclusion, lead VHH E4-1 and its optimized variant VHH Z70 have both the capacity to inhibit the aggregation of Tau *in vitro* and their relative activity is related to their affinity for Tau.

**Fig 3.**
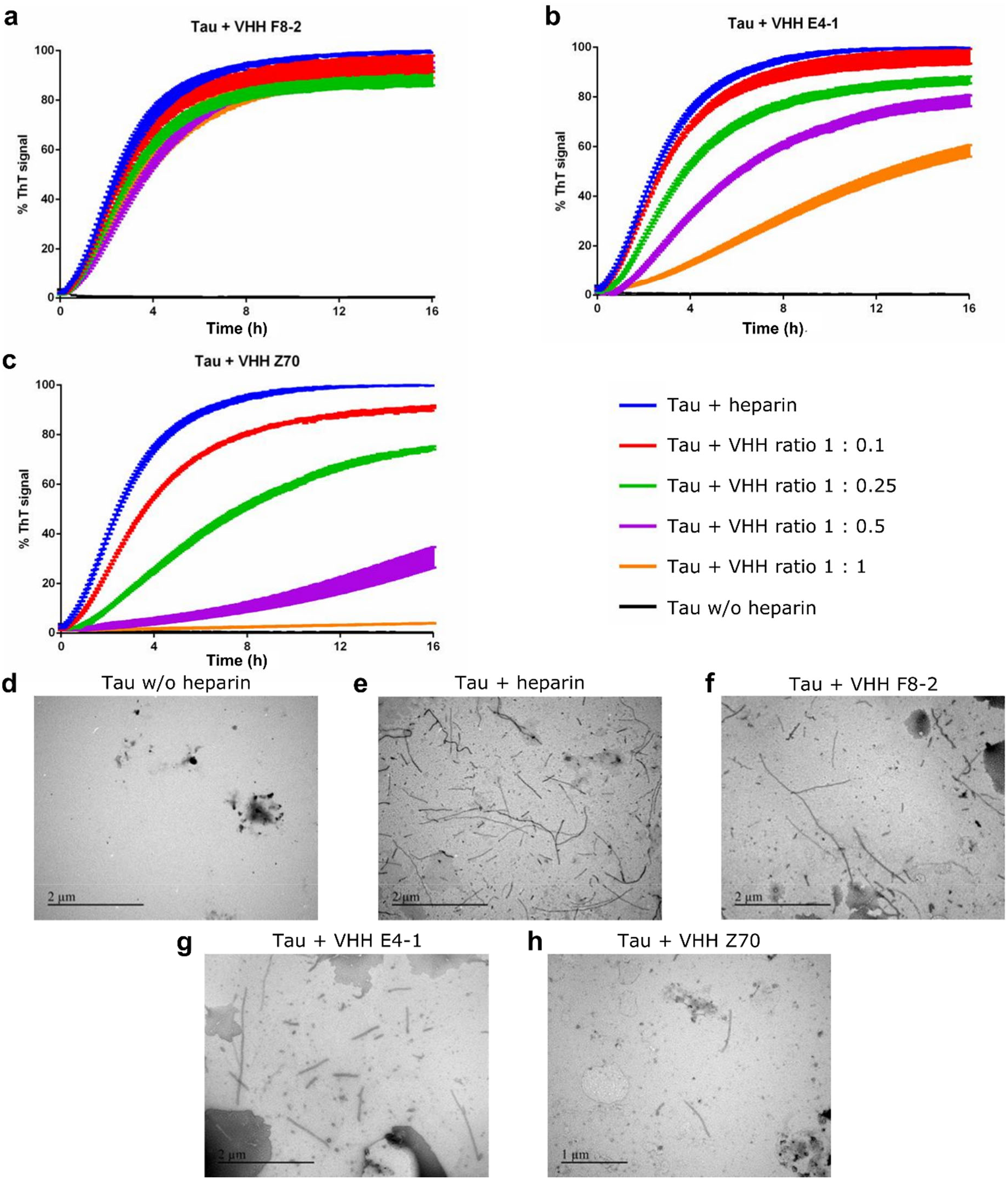
VHH E4-1 and VHH Z70 inhibit *in vitro* Tau aggregation Aggregation of Tau (10 µM) in the absence of heparin (black curve), in the presence of heparin and of increasing concentration of **a** VHH F8-2 **b** VHH E4-1 and **c** VHH Z70 (0, 1, 2.5, 5 and 10 µM) monitored by Thioflavin T fluorescence at 490 nm (n=3). Error bars: s.e.m, **d-h** Transmission electron microscopy images at the end point of the aggregation assays **d** in the absence of heparin or **e** in the presence of heparin and **f-h** in the presence of heparin and the additional presence of **f** VHH F8-2 **g** VHH E4-1 **h** VHH Z70 (for Tau/VHH molar ratio of 1 : 1) (n=2).

### Inhibition of Tau seeding in HEK293 Tau repeat domain (RD) P301S FRET Biosensor seeding reporter cells

The capacity of VHH E4-1 and VHH Z70 to block the seeding in the HEK293 Tau RD P301S FRET Biosensor reporter cell line model was next investigated. This cell line constitutively expresses Tau RD (MTBD), with a P301S mutation, fused to either CFP (Cyan Fluorescent Protein) or YFP (Yellow Fluorescent Protein) that together generate a FRET (Forster Resonance Energy Transfer) signal upon MTBD-P301S aggregation seeding ^49^. For cells treated with HEPES buffer only, FRET signal is detected neither by confocal microscopy nor by flow cytometry (**Fig. 4a-b-e**). The intracellular aggregation of MTBD-P301S protein is induced by treating the cells with Tau seeds, the MTBD fragment *in vitro* aggregated in HEPES buffer with heparin, associated to liposomes to help cell penetration^49^, leading to a FRET signal (yellow fluorescence by confocal and 16% ± 0.8% FRET-gated positive cells by flow cytometry, **Fig. 4c-d-e**). In addition, VHH F8-2 was transfected one day prior to MTBD seed treatment as negative control since its binding is outside the MTBD and is thus expected to not affect the seeding in the reporter cells (15.4% ± 1% FRET-gated positive cells, **Fig. 4 e-g**). The mCherry-VHH fusion proteins allowed to visualize transfected cells (red-orange fluorescence by confocal, **Fig 4 f-g**) and to detect FRET signal selectively in mCherry-VHH positive cells (mCherry-gated and FRET-gated positive cells by flow cytometry, **Fig. 4 h-j**). The FRET signal reduction for VHH E4-1 positive cells was not significant, with a percentage of FRET decreasing to 15.6 ± 0.9%, compare to 18.5 ± 1.3% FRET signal for VHH F8-2 negative control (15.7% seeding inhibition, **Fig. 4h-j**). Conversely, VHH Z70 clearly affected the intracellular seeding of MTBD-P301S aggregation, as the observed FRET signal for the corresponding transfected cells was significantly decreased to 10.9 ± 0.7% (41% seeding inhibition, **Fig. 4h-j**). Based on these measurements, we concluded that the amount of intracellular aggregates of MTBD-P301S Tau was reduced by more than 40% in the presence of the VHH Z70, showing the efficiency of VHH Z70 to block Tau seeding in this cellular model.

**Fig 4.**
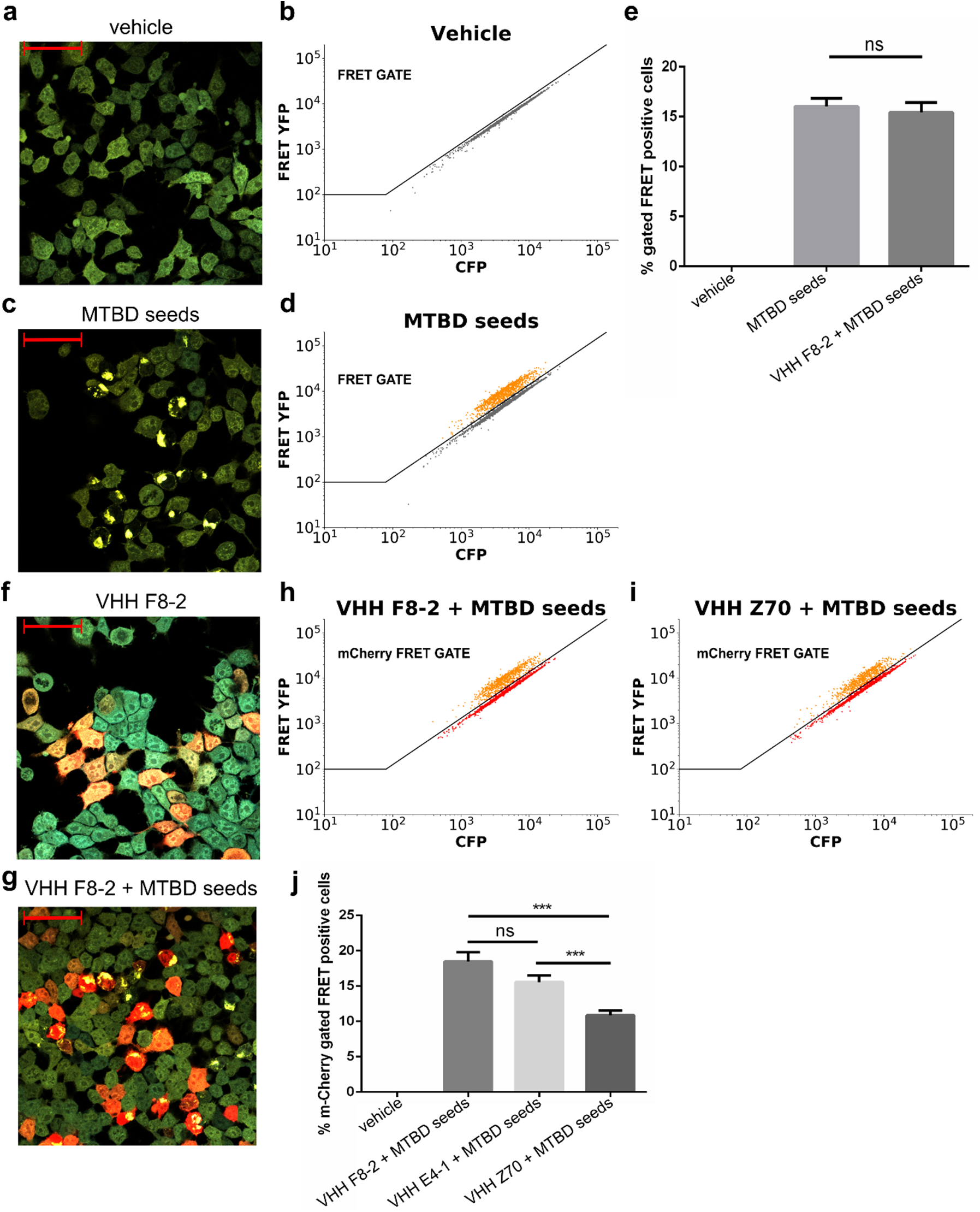
VHH Z70 blocks intracellular seeding of Tau MTBD in HEK293 biosensor cells. **a-d** Analysis of Tau seeding in HEK293 Tau RD P301S FRET Biosensor cells **a** by confocal microscopy for cells transfected with vehicle (HEPES buffer) **b** by flow cytometry with a FRET-gate for cells transfected with vehicle **c** by confocal microscopy for cells transfected with MTBD seeds. Positive cells that have incorporated MTBD seeds show yellow dots corresponding to FRET signal **d** by flow cytometry with a FRET-gate for cells transfected with MTBD seeds **e** Percentage of FRET positive cells determined from flow cytometry data for cells transfected as in **b, d** or transfected with VHH F8-2 followed by MTBD seeds. **f-i** Analysis of Tau seeding in HEK293 Tau RD P301S FRET Biosensor cells **f-g** by confocal microscopy for cells transfected with **f** VHH F8-2 or **g** with VHH F8-2 followed by MTBD seeds. Cells transfected with VHHs have a orange red color from the mCherry reporter, FRET is visualized as yellow dots. **h-i** by flow cytometry with a mCherry FRET gate for cells transfected with **h** VHH F8-2 followed by MTBD seeds (n=3) or **i** VHH Z70 followed by MTBD seeds (n=3) **j** Percentage of mCherry-gated FRET positive cells, determined from flow cytometry data for cells as in **h, i** or for cells transfected with VHH E4-1 followed by MTBD seeds (n=3). A significant decrease of FRET signal, reporting a decrease intracellular MTBD aggregation, is observed in the presence of VHH Z70. Error bars: s.e.m ***p < 0.001, Mann-Whitney U-test. Error bars: s.e.m.

### Inhibition of Tau seeding in the Tg tauopathy mouse model THY-tau 30

The next step was to test the capacity of VHH Z70 to block Tau seeding in a tauopathy mouse model. This model consists in the intra-cerebro cranial injection of human AD (h-AD) brain-derived material to robustly induce Tau pathology on a relatively short time scale (1 month), in 1-month-old Tg THY-tau30 mice^24,50^. At 1-month-old, these mice have low endogenous Tau pathology, which allowed to evaluate the seeding activity associated to the injected human-derived materials.

To assay the capacity of VHH Z70 to mitigate the tauopathy in this model, VHH Z70 was expressed as a fusion protein with mCherry inside brain cells, following infection by LVs (**Supplementary Fig. 8**). LVs were delivered using intra-cerebrocranial injection in each brain hemisphere, 2 weeks prior to the stereotaxic h-AD seed delivery at the same coordinates in the hippocampus region (**Fig. 5a**). Control experiments consisted in injection of PBS buffer instead of the h-AD brain extract and a VHH directed against GFP (VHH anti-GFP), which is not present in mouse brains, instead of VHH Z70. The region of expression of VHH Z70 in the brain was monitored in both hemispheres using the mCherry fusion as a reporter. The level of Tau pathology was evaluated, 1-month post-injection of the seeds, by immunohistochemistry with the monoclonal AT8 antibody. AT8 binds pS202:pT205:pS208 that is present in the PHFs and other abnormaly phosphorylated forms of Tau^51^, and is widely used as an AD-specific antibody^52^. AT8 Tau epitope detection was quantified between bregma −2.06 to −2.92 for a surface area where the pathology is seeded as determined in the positive control group injected with h-AD brain extract and treated with VHH anti-GFP. AT8 immunoreactivity indeed showed a significant difference between the pathology in this positive control group and the negative control group injected with PBS and treated with VHH anti-GFP (**Fig. 5 b-c, Supplementary Fig. 9 a-b**). In the animals injected with h-AD brain extract and expressing VHH Z70, a significant decrease of the average AT8 labeling compared to the positive control group is observed (**Figure 5 b-c, Supplementary Fig. 9 b-c**). In this group, the expression of VHH Z70 between bregma −2.06 and −2.92 was monitored using mCherry immunoreactivity (**Supplementary Fig. 10**). The decrease detected in the quantified area showed the positive effect of VHH Z70 on mitigating the seeded pathology. We concluded that intracellular immunization with VHH Z70 prevents the seeding induced by injection of extracellular h-AD brain extract in THY-tau30.

**Figure 5.**
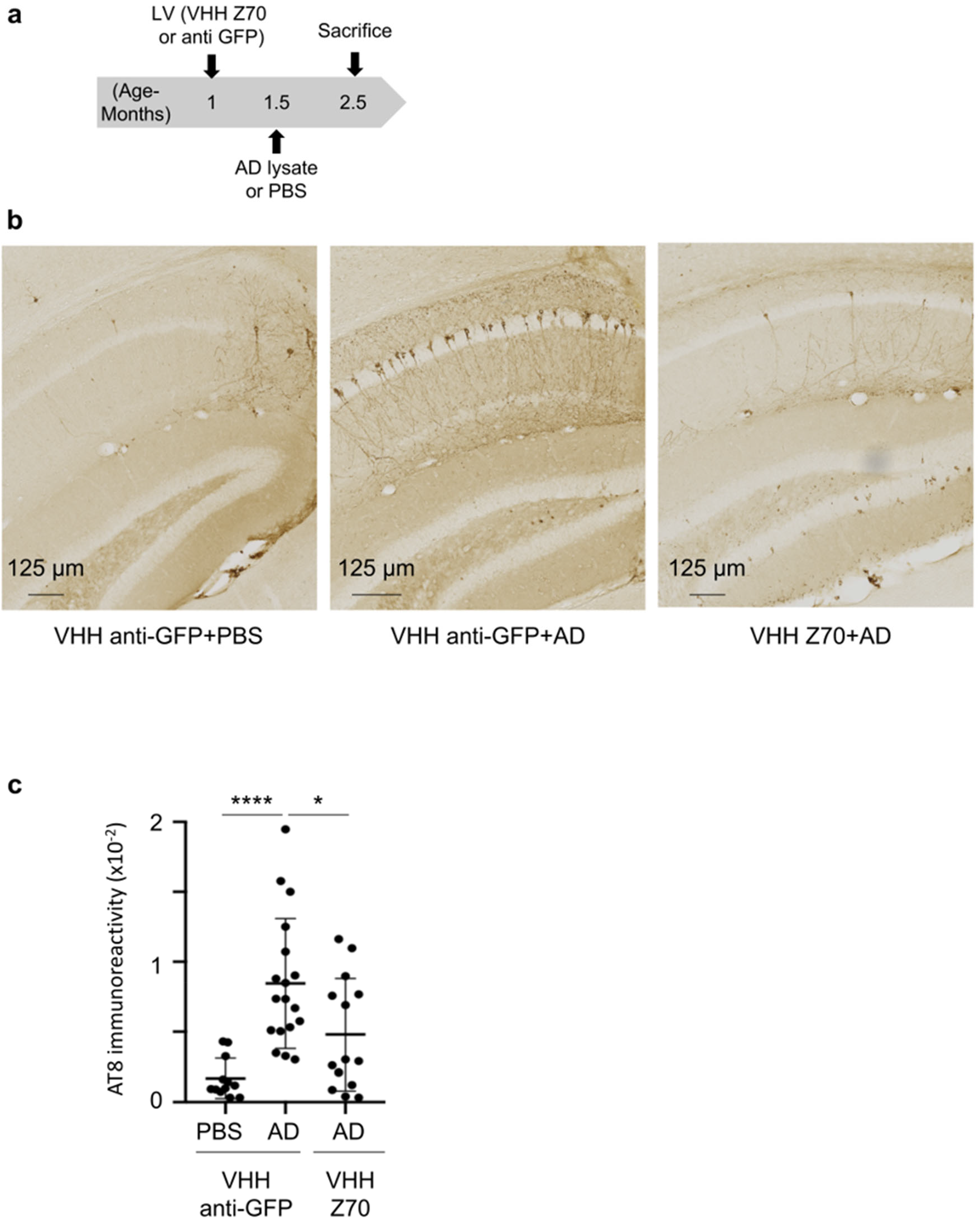
VHH Z70 reduces human Tau seeding activity induced by h-AD brain-derived material in THY-tau30 Tg mouse model. **a**. Bilateral intracranial injection of LVs to express either VHH anti-GFP or VHH Z70 was performed in one-month old THY-tau30 mice. Induction of the tauopathy occurred by a second bilateral injection, at the same coordinates, fifteen days after with human h-AD or PBS buffer. Mice were sacrificed and perfused one-month latter (aged 2.5 months). **b**. The whole brain were processed for immunohistochemical analysis using AT8. Three groups were analyzed corresponding to : injection of LVs VHH anti-GFP followed by PBS injection (LVs VHH anti-GFP + PBS; n=6), of LVs VHH anti-GFP followed by h-AD injection (LVs VHH anti-GFP+ h-AD; n=9) and of LVs VHH Z70 followed by h-AD injection (LVs VHH Z70 + h-AD; n=7). Enlarged images are taken from the hippocampus (injection site, AP : −2.46). Scale bars are indicated on the figure. **c**. Each data point corresponds to the quantification for one hemisphere. Results are presented as AT8 immunoreactivity (immunoreactive area normalized to the whole area). Error bars correspond to the S.D., **** p < 0.0001, * p < 0.5, Mann-Whitney U-test.

## Discussion

Tau immunotherapy is an attractive strategy in tauopathies to bind to and to clear extracellular and/or intracellular pathological species of the Tau protein to slow disease progression. Indeed, by targeting different Tau epitopes, immunotherapy studies showed a reduction of Tau pathology and cognitive deficit in different mouse models of tauopathy^18–25^.

We succeeded in selecting a VHH targeted to a specific region in a rather large disordered protein, in one single screen. NMR epitope mapping was particularly helpful to select for VHHs binding the MTBD. However, NMR epitope mapping does not allow for the unambiguous identification of the sole residues directly involved in the interaction. The decrease in NMR resonance intensity (**Fig. 1**, Supplementary **Fig 1-4, 7**) can result from local immobilization of the disordered protein due to the binding, decreasing local tumbling and thus increasing relaxation. Accordingly, the Tau domain involved in the VHH interaction, which contained the PHF6 and PHF6*, was described as presenting local extended secondary structure^45^ and thus represents a relatively rigid stretch that could explain the extended region with resonance broadening upon binding. Alternatively, decrease resonance intensity can be due to chemical exchange between bound and unbound states that can result in line broadening, depending on the affinity and chemical shift change resulting from the interaction. In this case, the observed binding in the repeat region of Tau, given the level of sequence redundancy, could correspond to binding to R2 or R3 repeats, even if one is a secondary site of lower affinity to which NMR remained sensitive. However, alternative methodology by yeast two-hybrid has allowed for the identification of a single binding site (_305_SVQIVYKPV_313_), in agreement with the SPR data that are compatible with a one-to-one binding model.

The main interest is that VHH Z70 binds to the PHF6 in the MTBD, while the majority of epitopes targeted in clinical trial are located in the N- or C-terminal regions^26^. Some studies show that targeting N-terminal species could block Tau uptake and its transfer between neurons^53^, and could reduce accumulation of Tau in the brain of mouse model of tauopathy^54^. Nevertheless, antibodies targeting N- and C-terminal epitopes bring the risk of binding Tau sequences eliminated by proteolysis. They are also unlikely to interfere with aggregation and seeding, other crucial aspects of the pathology. Conversely, Bepranemab, a humanized monoclonal IgG4 antibody binding to the central region of Tau (235–246), near the MTBD, was reported to inhibit seeding in a cellular assay and a mouse model^24,55^. Similarly, treatment with a humanized IgG antibody, binding the R2 and R4 repeats within the MTBD, decreases the level of sarkosyl-insoluble Tau in brain of a mouse model of Tau seeding^56^. VHH Z70 has similar properties but by acting directly intracellularly.

Here, we demonstrated the interest of targeting the Tau PHF6 motif, which participates to the aggregation process^11^ and which is found in the core of Tau fibrils in AD^14–17^. In addition, although the mechanisms leading to the aggregated Tau species is not fully understood, accessibility of the PHF6*/ PHF6 peptides is a molecular feature identified as involved in the aggregation process. The residues from these peptides are proposed to be shielded in a dynamic hairpin conformations in native Tau, while exposed PHF6 residues would increase Tau sensitivity to aggregation^57^. The conversion from an inert Tau monomer to a seed-competent monomer would thus involve an increased accessibility of the PHF motifs^57^. Interestingly, several chaperons with Tau anti-aggregation activities, such as Hsc70, Hsp60 and DnaJA2, bind to regions overlapping PHF6, and in a weaker manner PHF6*. Thus, one major mechanism of the anti-aggregation activities of these chaperons is likely the binding of Tau in the PHF region^58^.

Additionally, the properties of this initial VHH were improved, such that the optimized VHH Z70 variant of the lead VHH E4-1 demonstrated better inhibitory activity both in *in vitro* aggregation assays and in a cellular model of seeding, blocking the recruitment of the intracellular Tau MTBD. The affinity reached the 100nM range and remained an important parameter to optimize as we observed a higher anti-aggregation activity for VHH Z70 compared to VHH-E4-1 in the *in vitro* assay (**Fig. 3**). The poorer seeding inhibition capacity of VHH-E4-1 is likely due in addition to its poor intracellular activity compared to VHH Z70 (**Fig. 4**).

Accordingly, VHH Z70 decreases the Tau pathology in an established mouse model of tauopathy^24^. In this injection model, VHH Z70 mechanism of action likely results from blocking Tau seeding, limiting the accumulation of pathological Tau. According to our *in vitro* results, nucleation is probably blocked by the binding of the VHHZ70 to monomeric Tau, preventing its recruitment by the seeds. We cannot however exclude the possibility that VHH Z70 also binds to the seeds, given the lack of a precise definition of the nature of these seeds: seed-competent monomeric, oligomeric or fibrillar Tau.

VHHs have entered the real world of therapeutics^59^, and as they can be delivered as genes, the synergy with the progress in viral vector-mediated gene delivery could open the way for feasible treatments of tauopathies.

## Materials and methods

### Screening and Selection of VHHs directed against Tau protein

The Nali-H1 library of VHHs was screened against the recombinant biotinylated-Tau as described previously^43,48^. Non-absorbed Phage ELISA assay using avidin-plates and biotinylated-Tau Antigen (5µg/ml) was used for cross-validation of the selected clones^60^.

### Production and purification of VHHs

Competent *Escherichia coli* BL21 (DE3) bacterial cells were transformed with the various PHEN2-VHH constructs. Recombinant *E. coli* cells produced proteins targeted to the periplasm after induction by 1 mM IPTG (isopropylthiogalactoside). Production was pursued for 4 hours at 28°C before centrifugation to collect the cell pellet. Pellet was suspended in 200 mM Tris-HCl, 500 mM sucrose, 0.5 mM EDTA, pH 8 and incubated 30 min on ice. 50 mM Tris-HCl, 125 mM sucrose, 0,125 mM EDTA, pH 8 and complete protease inhibitor (Roche) were then added to the cells suspension and incubation continued 30 min on ice. After centrifugation, the supernatant, corresponding to the periplasmic extract, was recovered. The VHHs were purified by immobilized-metal affinity chromatography (HisTrap HP, 1mL, Cytiva) followed by size exclusion chromatography (Hiload 16/60, Superdex 75, prep grade, Cytiva) in NMR buffer (50 mM sodium phosphate buffer (NaPi) pH 6.7, 30 mM NaCl, 2.5 mM EDTA, 1 mM DTT).

### Production and purification of labelled ^15^N Tau 2N4R, ^15^N Tau 2N3R and ^15^N Tau MTBD

pET15b-Tau recombinant T7lac expression plasmid was transformed into competent *E. coli* BL21 (DE3) bacterial cells. A small-scale culture was grown in LB medium at 37 °C and was added at 1:10 V/V to 1L of a modified M9 medium containing MEM vitamin mix 1X (Sigma-Aldrich), 4 g of glucose, 1 g of ^15^N-NH_4_Cl (Sigma-Aldrich), 0.5 g of ^15^N-enriched Isogro (Sigma-Aldrich), 0.1 mM CaCl_2_ and 2 mM MgSO_4_. Recombinant ^15^N Tau production was induced with 0.5 mM IPTG when the culture reached an optical density at 600nm of 0.8. Proteins were first purified by heating the bacterial extract, obtained in 50 mM NaPi pH 6.5, 2.5 mM EDTA and supplemented with complete protease inhibitors cocktail (Sigma-Aldrich), 15 min at 75 °C. The resulting supernatant was next passed on a cation exchange chromatography column (Hitrap SP sepharose FF, 5mL, Cytiva) with 50 mM NaPi pH 6.5 and eluted with a NaCl gradient. Tau proteins were buffer-exchanged against 50 mM ammonium bicarbonate (Hiload 16/60 desalting column, Cytiva) for lyophilization. The same protocol was used to produce and purify Tau 2N3R isoform and Tau[245-368] (designated MTBD, also called K18 fragment). Detailed procedure can be found in ^61^.

### Production and purification of SUMO-Tau peptides

cDNA encoding peptides Tau[273-318], was amplified from Tau 2N4R cDNA by PCR. cDNA was cloned by a ligation independent protocol into vector pETNKI-HisSUMO3-LIC as described^62^. Tau peptide was expressed as N-terminal SUMO protein fusion with a N-terminal HisTag. His-SUMO-Tau peptide was purified by affinity chromatography on Ni-NTA resin followed by size exclusion chromatography (Hiload 16/60, Superdex 75, prep grade, Cytiva) in SPR buffer (HBS-EP+, GE Healthcare).

### Nuclear Magnetic Resonance Spectroscopy Experiments

Analysis of the ^15^N Tau/VHH interactions were performed at 298K on a Bruker Avance Neo 900MHz spectrometer equipped with cryogenic probe. TMSP (trimethyl silyl propionate) was used as internal reference. Lyophilized ^15^N Tau were diluted in a buffer containing 50 mM NaPi, 30 mM NaCl, 2.5 mM EDTA, 1 mM DTT, and 10% D_2_O, pH 6.7 and mixed with VHH at 100 μM final concentration for each protein. 200 μL of each mix in 3 mm tubes were sufficient to obtain the 2D ^1^H, ^15^N HSQC spectra with 32 scans. ^1^H, ^15^N HSQC were acquired with 3072 and 416 points in the direct and indirect dimensions, for 12.6 and 25 ppm spectral windows, in the ^1^H and ^15^N dimensions respectively. Data were processed with Bruker Topspin 3.6 and analyzed with Sparky (T. D. Goddard and D. G. Kneller, SPARKY 3, University of California, San Francisco).

### Optimization of VHH E4-1 for intracellular expression

VHH E4-1 was amplified from pHEN2 plasmid (oligonucleotides 3390 and 3880 in **Supplementary Fig. 11**) using Taq polymerase with 14 mM MgCl_2_ and 0.2 mM MnCl_2_ and a modified nucleotide pool^63^. The amplified cDNAs were transformed in yeast Y187 strain, together with a digested empty derivative of pGADGH vector^64^, allowing recombination by gap repair in the vector. The VHH cDNAs are expressed as preys, with a C-terminal Gal4-activation domain fusion (E4-1-Gal4AD). A library of 2.1 million clones was obtained, collected and aliquoted. Tau variant 0N4R isoform (NM_016834.4) was expressed as bait with a C-terminal fusion with lexA (Tau-LexA) from pB29 vector, which is derived from the original pBTM116^65^. The library was screened at saturation, with 20 million tested diploids, using cell-to-cell mating protocol ^66^. A single clone was selected, named VHH Z70. A one-to-one mating assay was used to test for interaction using a mating protocol with L40ΔGal4 (mata) transformed with the bait and Y187 (matα) yeast strains transformed with the prey ^66^. The interaction pairs were tested in triplicate on selective media by streak.

### Tau fragment library construction

Tau cDNA (NM_016834.4) was amplified from Tau-LexA bait vector (oligonucleotides 6690 and 6972 **Supplementary Fig. S11**). 5 µg of the PCR product was subjected to Fragmentase® treatment (New England Biolab, NEB) until a smear of fragments was detected around 400-500pb by agarose gel electrophoresis. The DNA fragments were purified by phenol/chloroform extraction and ethanol precipitation. The DNA fragments were next subjected to end repair (NEB) and dA-tailing adaptation, using Blunt/TA ligase master mix with NEBNext® Adaptor hairpin loop (NEB), followed by AMPure XP bead (Beckman Coulter) purification. After USER® enzyme digestion (NEB), DNA fragments were amplified (oligonucleotides 10829 and 10830 in **Supplementary Fig. 11**) with 15 cycles of PCR using NEBNext® Q5® Hot Start HiFi PCR Master Mix (NEB), which allowed to add Gap Repair recombination sequences for the cloning in Gal4-AD prey plasmid pP7. The library comprised 50000 independent clones.

### Tau fragment library screening

The coding sequence for VHH Z70 was PCR-amplified and cloned into pB27 as a C-terminal fusion to LexA (LexA-VHHZ70). The construct was used to produce a bait to screen the Tau fragments library constructed into pP7. pB27 and pP7 derived from the original pBTM116 ^65^ and pGADGH ^64^ plasmids, respectively. The Tau fragment library was screened using a mating approach with YHGX13 (Y187 ade2-101::loxP-kanMX-loxP, matα) and L40ΔGal4 (mata) yeast strains^66^. 90 His+ colonies corresponding to 267.10^3^ tested diploids were selected on a medium lacking tryptophan, leucine and histidine. The prey fragments of the positive clones were amplified by PCR and sequenced at their 5’ and 3’ junctions.

### Surface Plasmon Resonance experiments

Affinity measurements were performed on a BIAcore T200 optical biosensor instrument (Cytiva). Recombinant Tau proteins were biotinylated with 5 molar excess of NHS-biotin conjugates (Thermofisher) during 4 hours at 4 °C. Capture of biotinylated Tau was performed on a streptavidin SA sensorchip in HBS-EP+ buffer (Cytiva). One flow cell was used as a reference to evaluate nonspecific binding and provide background correction. Biotinylated-Tau was injected at a flow-rate of 30 μL/min, until the total amount of captured Tau reached 500 resonance units (RUs). VHHs were injected sequentially with increasing concentrations ranging between 0.125 and 2 µM in a single cycle, with regeneration (3 successive washes of 1M NaCl) between each VHH. On the other hand, VHH Z70 was immobilized on a CM5 chip in HBS-EP+ buffer (Cytiva) and increasing concentrations, ranging between 0.125 and 2 µM of the SUMO-Tau peptide, were successively injected. Single-Cycle Kinetics (SCK) analysis^52^ was performed to determine association *k*_on_ and dissociation *K*_off_ rate constants by curve fitting of the sensorgrams using the 1:1 Langmuir model of interaction of the BIAevaluation sotware 2.0 (Cytiva). Dissociation equilibrium constants (Kd) were calculated as *k*_on/_*K*_off_.

### *In vitro* kinetic aggregation assays

Tau 2N4R aggregation assays were performed with 10 μM Tau and with increasing concentrations of VHHs (between 0 and 10 μM) in buffer containing 50 mM MES pH 6.9, 30 mM NaCl, 2.5 mM EDTA, 0.3 mM freshly prepared DTT, 2.5 mM heparin H3 (Sigma-Aldrich) and 50 μM Thioflavin T (Sigma-Aldrich), at 37°C. Experiments were reproduced 3 times in triplicates for each condition. The resulting fluorescence of Thioflavin T was recorded every 5 min/cycle within 200 cycles using PHERAstar microplate-reader (BMG labtech. The measures were normalized in fluorescence percentage, 100% being defined as the maximum value reached in the positive Tau control, in each experiment.

### Transmission Electron Microscopy

The same samples from the *in vitro* aggregation assays were recovered and a 10 µl sample of Tau or Tau:VHH ratio 1:1 condition was loaded on a formvar/carbon-coated grid (for 5 min and rinsed twice with water). After drying, the grids were stained with 1% uranyl acetate for 1 min. Tau fibrils were observed under a transmission electron microscope (EM 900 Zeiss equipped with a Gatan Orius 1000 camera).

### Seeding assays in HEK293 reporter cell-line

Stable HEK293 Tau RD P301S FRET Biosensor cells (ATCC CRL-3275) were plated at a density of 100k cells/well in 24-well plates. For confocal analysis, cells were plated on glass slides coated with poly-D-lysine and laminin at a density of 100k cells/well in 24-well plates. At 60% confluency, cells were first transiently transfected with the various pmCherry-N1 plasmid constructs allowing expression of the mCherry-fused VHHs. Transfection complexes were obtained by mixing 500 ng of plasmid diluted in 40 µl of opti-MEM medium, which included 18.5 µL (46.25% v/v) of opti-MEM medium with 1.5 µL (3.75% v/v) Lipofectamine 2000 (Invitrogen). Resulting liposomes were incubated at room temperature for 20 min before addition to the cells. Cells were incubated for 24 hours with the liposomes and 1 ml/well of high glucose DMEM medium (ATCC) with Fetal Bovine Serum 1% (Life technologies). The transfection efficiency was estimated to reach about 46%, for all mCherry-fused VHH plasmids (**Supplementary Fig. 12**). Eight µM of recombinant MTBD seeds were prepared *in vitro*, in the presence of 8 µM heparin, as described^49^. Cells were then treated with MTBD seeds (10 nM/well) in the presence of transfection reagents forming liposomes as here above described.

### Confocal analysis

Cells were first washed twice with PBS and fixed in 4% paraformaldehyde (PFA) for 20 min and next washed 3 times with 50 mM NH_4_Cl in PBS. Glass slides were mounted with DAKO mounting medium (Agilent). Fluorescence imaging acquisitions were performed using an inverted confocal microscope (LSM 710, Zeiss, Jena, Germany) with a 40-times oil-immersion lens (NA 1.3 with an optical resolution of 176 nm). A focal plane was collected for each specimen. Images were processed with ZEN software.

### FRET Flow Cytometry

Cells were recovered with trypsin 0,05% and fixed in 2% PFA for 10 min, then suspended in PBS. Flow cytometry was performed on an ARIA SORP BD (acquisition software FACS DIVA V7.0 BD, Biosciences). To measure CFP emission fluorescence and FRET, cells were excited with a 405 nm laser. The fluorescence was captured with either a 466/40 or a 529/30 nm filter, respectively. To measure YFP fluorescence, a 488 nm laser was used for excitation and emission fluorescence was captured with a 529/30 nm filter. mCherry cells were excited with a 561 nm laser and fluorescence was captured with a 610/20 nm filter. To selectively detect and quantify FRET, gating was used as described ^49,68^. The FRET data were quantified using the KALUZA software analyze v2. Three independent experiments were done in triplicate or quadruplicate, with at least 10,000 cells per replicate analyzed.

### Animals

The study was performed in accordance with the ethical standards as laid down in the 1964 Declaration of Helsinki and its later amendments or comparable ethical standards. The experimental research has been performed with the approval of an ethical committee (agreement APAFIS#2264-2015101320441671 from CEEA75, Lille, France) and follows European guidelines for the use of animals. The animals (males and females) were housed in a temperature-controlled (20-22°C) room maintained on a 12 h day/night cycle with food and water provided *ad libitum* in specific pathogen free animal facility (n=5 mice per cage). Animals were allocated to experimental groups by randomization. AD brain extracts were obtained from the Lille Neurobank (fulfilling criteria of the French law on biological resources and declared to competent authority under the number DC-2008-642) with donor consent, data protection and ethical committee review. Samples were managed by the CRB/CIC1403 Biobank, BB-0033-00030.

### Stereotaxic injection of THY-tau30 Tg Mice

THY-tau30 Tg mice express human 1N4R tau protein with two pathogenic mutations (P301S and G272V) under the control of the neuron-specific Thy1.2 promoter^69,70^. 1-month-old anesthetized THY-tau30 mice were submitted to stereotaxic intra-cerebrocranial injections (400ng in 2µl at 250nl/min with a Hamilton glass syringe) at the coordinates posterior AP: −2.46, midline ML: −1 and vertical depth DV: −2.3 of both brain hemispheres with LVs expressing either VHH Z70 with a N-terminal mCherry fusion protein (VHH Z70) or a VHH directed against the green fluorescent protein (VHH anti-GFP).

2 weeks later, these mice were submitted to injections of human AD brain homogenate (h-AD, 15 µg in 2 µl) or PBS (2 µl) at the same coordinates of both hemispheres, as previously described in detail^24^. The h-AD seeds consisted in a mixture of two post-mortem human brain extracts from tissues of confirmed AD patients (frontal cortex area, Braak stage VI, Brodmann area 10). The combination of injections resulted in three groups of 6-9 mice per group. The mice were sacrificed after a month delay from the injection of the h-AD brain extract.

### Tissue processing, immunohistochemistry and Tau pathology quantification

THY-tau30 were deeply anesthetized and trans-cardially perfused with ice-cold 0.9% saline solution and subsequently with 4% PFA for 10 min. The brains were immediately removed, fixed overnight in 4% PFA, washed in PBS, placed in 20% sucrose for 16 h and frozen in isopentane until further use. Free-floating coronal sections (40 µm thickness) were obtained using a cryostat microtome.

Cryostat sections were next used for immunohistochemistry. Non-specific binding was blocked by using ‘Mouse in Mouse’ reagent (1:100 in PBS, Vector Laboratories). Brain slices were next incubated with the primary monoclonal antibody AT8 (1:500, Thermo MN1020) in PBS-0.2% Triton X-100, 16h at 4°C. Labelling was amplified by incubation with an anti-mouse biotinylated IgG (1:400 in PBS-0.2% TritonTM X-100, Vector) followed by the application of the avidin-biotin-HRP complex (ABC kit, 1:400 in PBS, Vector) prior to addition of diaminobenzidine tetrahydrochloride (DAB, Vector) in Tris-HCl 0.1 mol/l, pH 7.6, containing H_2_O_2_ for visualization. Brain sections were mounted, air-dried, steadily dehydrated in ethanol (30%, 70%, 95%, 100%), cleared in toluene and cover-slipped with VectaMount (Vector Laboratories). Mounted brain sections were analyzed using stereology software (Mercator image analysis system; Explora Nova, La Rochelle, France). Threshold was established manually to present a minimum background and remained constant throughout the analysis. The region defined as quantification zone is from bregmas −2.06 to −2.92 (based on the Mouse Atlas, George Paxinos and Keith B.J. Franklin, Second Edition, Academic Press).

### ImmunoHistoFluorescence

Brain sections from mice injected with the LVs VHH Z70 with a N-terminal fusion to mCherry were saturated in normal goat serum (1/100, Vector), then were incubated with the primary polyclonal antibody anti-RFP targeting mCherry protein (1:1000, rabbit, Polyclonal, Rockland) 16h at 4°C in PBS-0.2% TritonTM X-100. Labelling was detected using a secondary anti-rabbit antibody (1:1000, Invitrogen) labelled with Alexa 488 for fluorescence detection (red). Section imaging was performed by microscopy using a slide scanner (Axioscan Z1-Zeiss) with a 20X objective.

### Statistical analysis

Data are presented as the means ± s.e.m for *in vitro* aggregation assays and reporter-cell seeding assays in HEK293 reporter cells. Experiments were performed at least in triplicate and obtained from three independent experiments. The Mann-Whitney U-Test was used for *in vivo* experiments to determine the p-value. The differences were considered significant at * p value < 0.05, **p value < 0.01, ***p < 0.001 and **** p value < 0.0001. Statistical analyses were performed with GraphPad Prism 5.01

## Acknowledgements

We thank Dr Z. Lens and Dr M. Aumercier for their help on the T200 biacore measurements. We also thank M. Tardivel and A. Bongiovanni for their help on the Zeiss confocal microscope, from the Photonic Microscopy Core BioImaging Center (BiCeL) and N. Jouy for the cytometry experiments, from the Flow Core Facility (BiCel).

## Author contribution statement

CD, ED, OZ, XH, MC, LB and IL performed, supervised the experiments and analyzed the data; F-XC performed and supervised the NMR experiments; JM prepared recombinant proteins; AA and J-CR performed the phage display VHH screen, yeast two-hybrid screens, the optimization of the VHHs and the epitope mapping; AL performed the EM experiments; SB prepared cells for confocal analysis and the LVs; RC supervised the animal experiments and performed the histological analysis; CD, ED and IL wrote the initial manuscript; J-CR, MC, XH and LB revised the manuscript, J-CR, LB and IL conceived the study.

## Abbreviations

AD: Alzheimer’s disease
CDR: complementarity-determining region
FR: framework domain
HSQC: heteronuclear single quantum correlation
IgG: Immunoglobulin G
LVs: lentiviral vectors
MTBD = RD: Tau microtubule binding domain = Tau repeat domain
NFTs: neurofibrillary tangles
NMR: nuclear magnetic resonance
PHF: paired helical filament
SPR: surface plasmon resonance
scFvs: single chain fragment of the variable domain of IgG
Tg: transgenic
VHH: Variable Heavy-chain of the Heavy-chain only antibody

