## Supplementary figures for "Inhibition of Tau seeding by targeting Tau nucleation core within neurons with a single domain antibody fragment"

### Supplementary Data

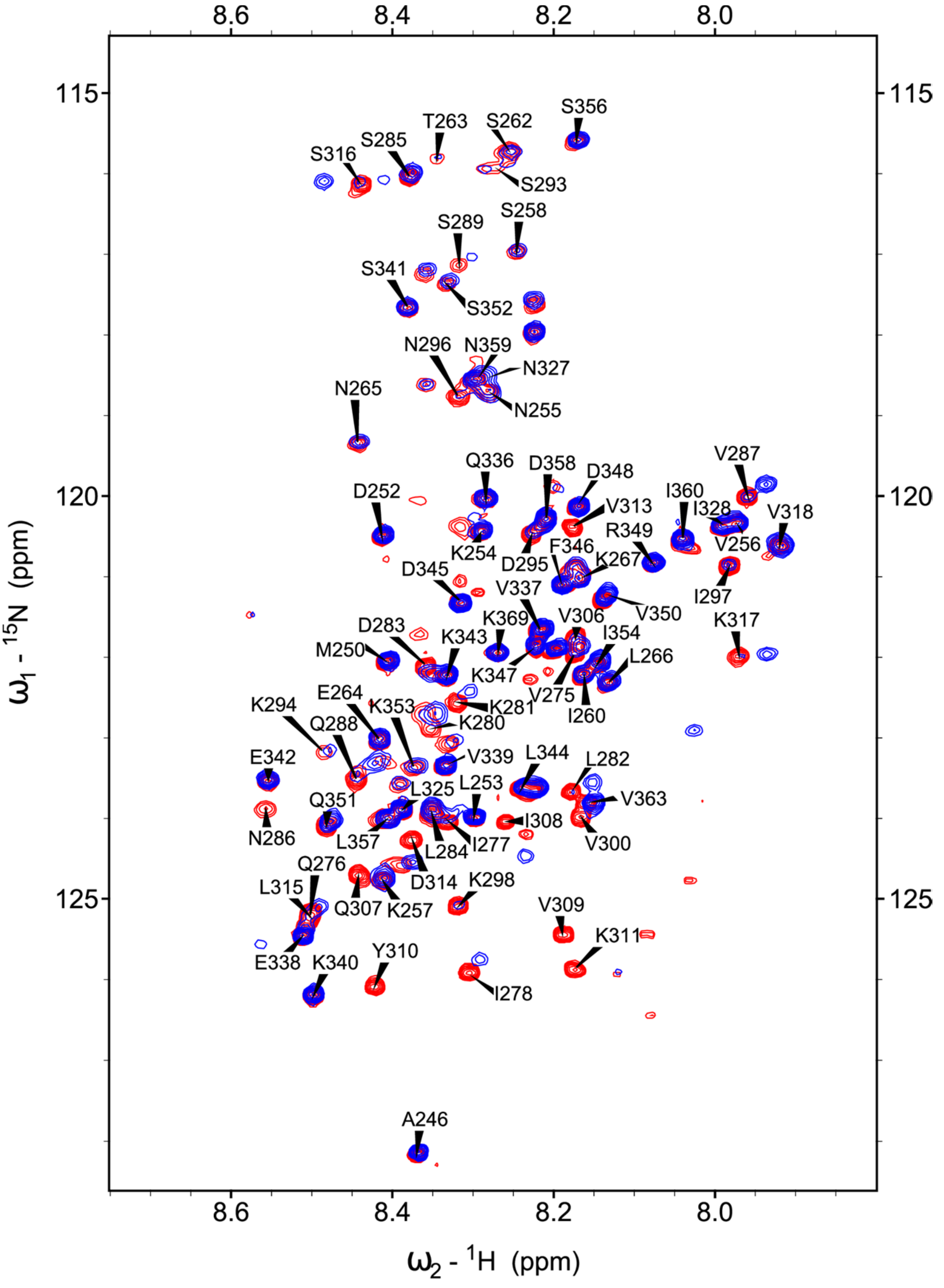


**Supplementary Figure 1 : Identification of VHH E4-1 epitope using Tau MTBD.** Overlay of $^{1}H$, $^{15}N$, HSQC two-dimensional spectra of $^{15}N$-labelled Tau MTBD (red) with $^{15}N$-labelled Tau MTBD mixed with non-labelled VHH E4-1 spectra (superimposed in blue).

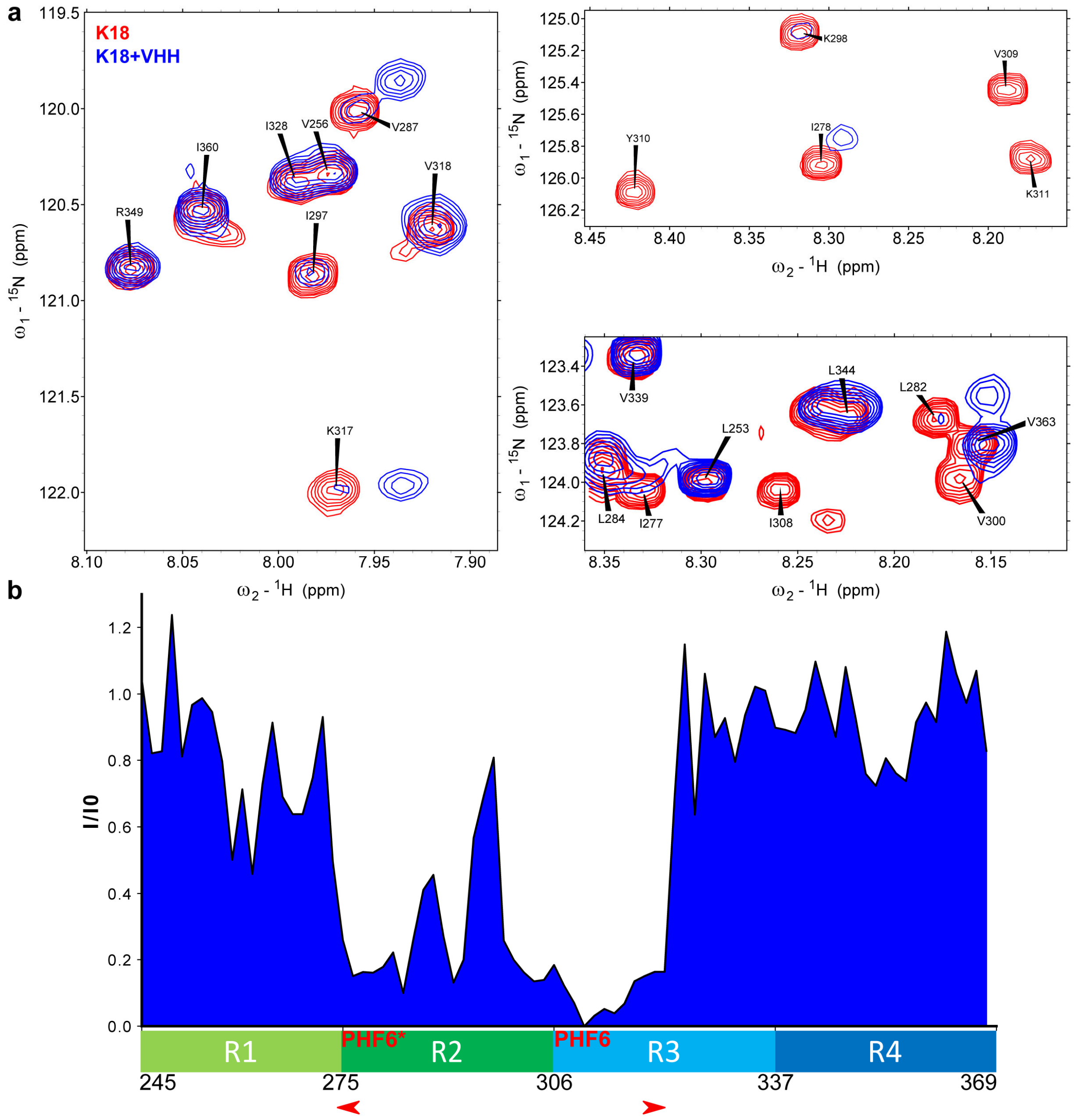


**Supplementary Figure 2 : Identification of VHH E4-1 epitope using Tau MTBD. a** : Overlays of $^{1}$H, $^{15}$N, HSQC enlargements of $^{15}$N-labelled Tau MTBD alone (red) or mixed with unlabelled VHH E4-1 (superimposed in blue). **b** : Intensity ratios I/I0 of corresponding resonances in the two-dimensional spectra of Tau MTBD with equimolar quantity of VHH E4-1 (I) or free in solution (I0) for residues along the Tau MTBD sequence. Overlapping resonances are not considered (x-axis is not scaled). The region containing the major broadened resonances corresponded to the R2-R3 repeats in the MTBD, as indicated by the red arrow heads. Localization of the PHF6* and PHF6 peptide sequences is indicated.

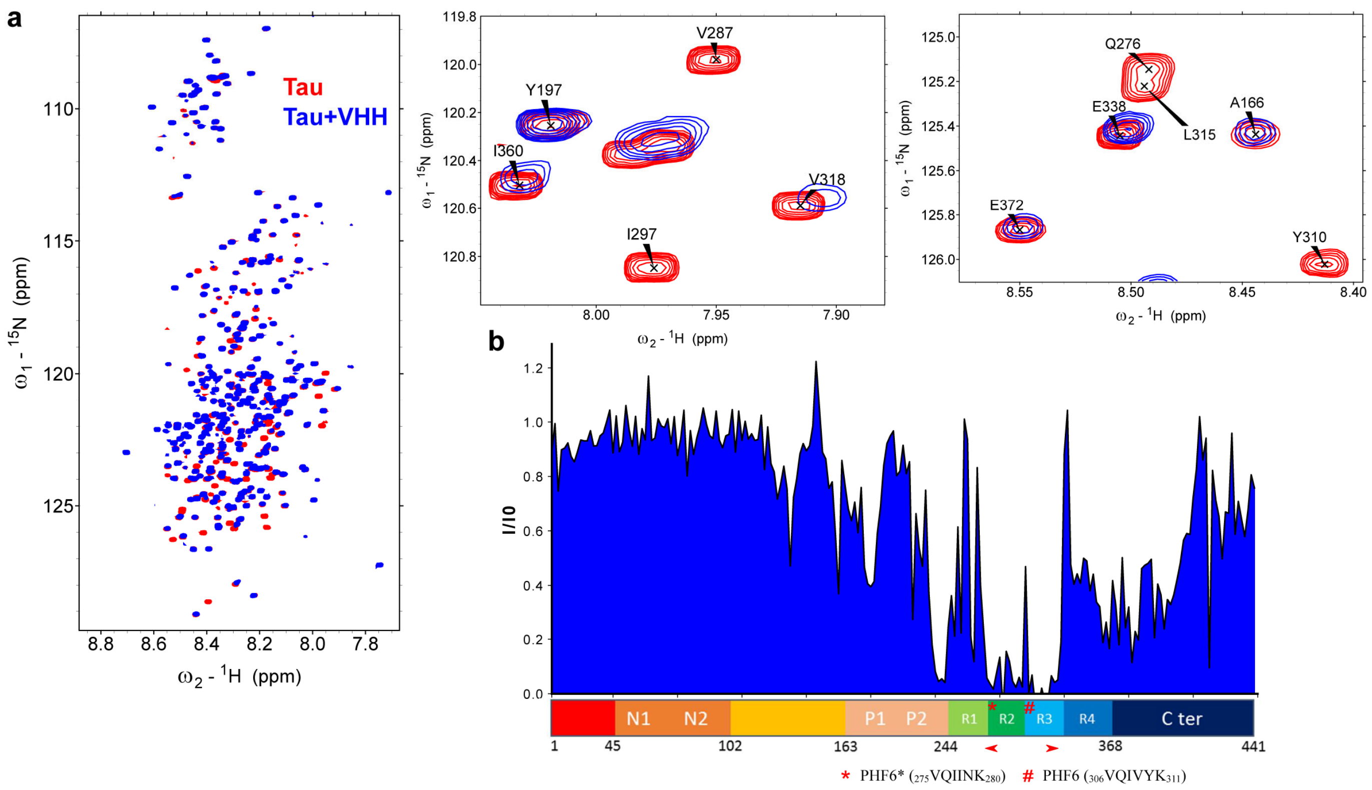


**Supplementary Figure 3 : Identification of VHH Z70 epitope using 2D HSQC NMR experiment. a** : Overlays of $^{1}$H, $^{15}$N, HSQC spectrum enlargements of $^{15}$N-labelled Tau, free (red) or mixed with non-labelled VHH Z70 (in blue). Spectra enlargements show broadened resonances corresponding to residues implicated in the interaction **b** : Intensities ratio I/I0 of corresponding resonances in the two-dimensional spectra of Tau with equimolar quantity of VHH Z70 (I) or free in solution (I0) for residues along the Tau sequence. Overlapping resonances are not considered (x-axis is not scaled). The red arrow heads indicate the region containing the corresponding major broadened resonances, which was mapped mostly on the R2-R3 repeats.

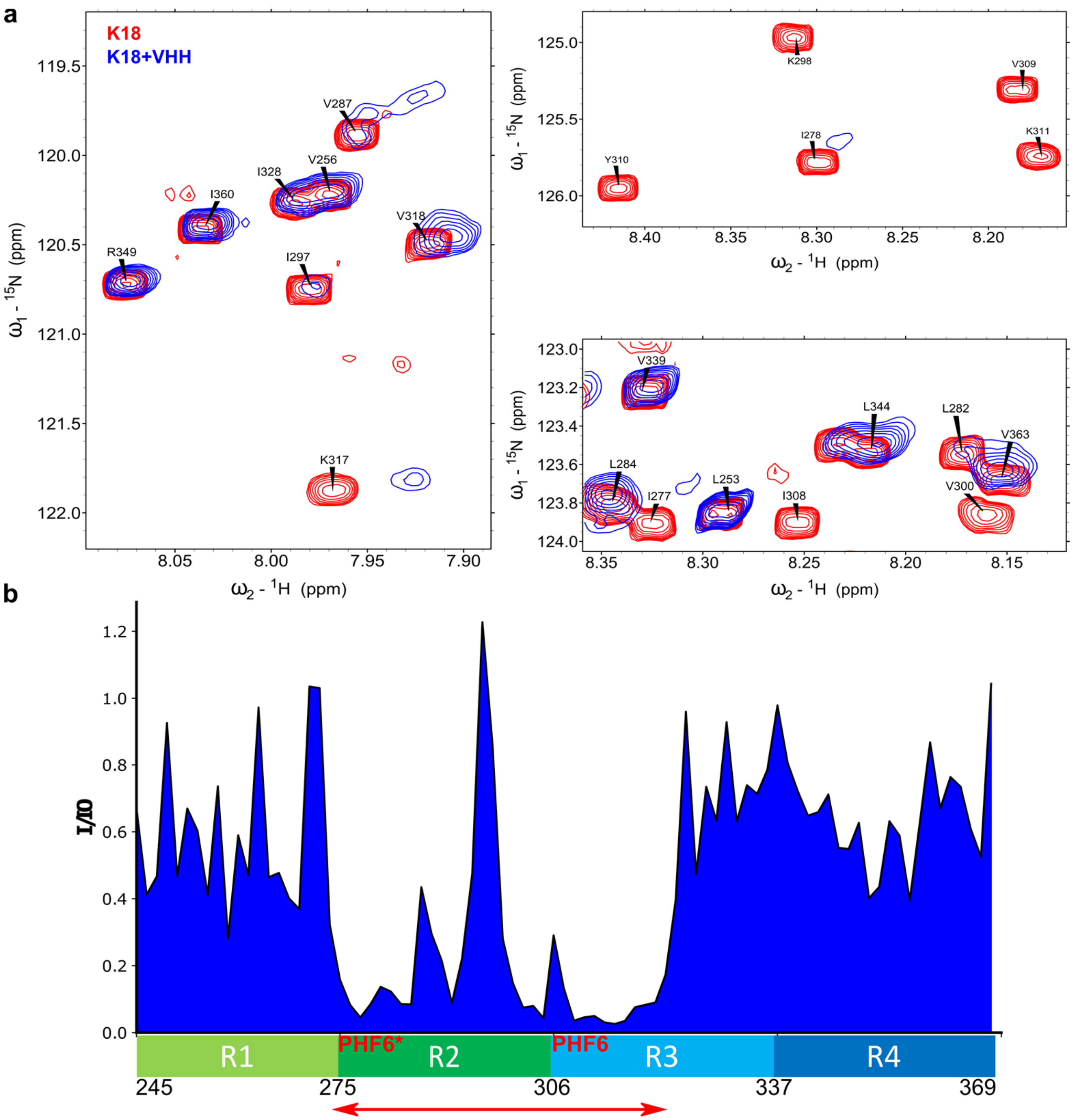


**Supplementary Figure 4 : Identification of VHH Z70 epitope using Tau MTBD. a** : Overlays of $^{1}$H, $^{15}$N, HSQC enlargements of $^{15}$N-labelled Tau MTBD alone (red) or mixed with VHH Z70 superimposed (in blue). **b** : Intensity ratios I/I0 of corresponding resonances in the two-dimensional spectra of Tau MTBD with equimolar quantity of VHH Z70 (I) or free in solution (I0) for residues along the Tau MTBD sequence. Overlapping resonances are not considered (x-axis is not scaled). The region containing the corresponding major broadened resonances mapped to the R2-R3 repeats in the MTBD. Localization of the PHF6* and PHF6 peptide sequences is indicated.

| VHH | $k_{on}$ (1/M.s) | $k_{off}$ (1/s) | Kd (nM) |
|---|---|---|---|
| E4-1 | 4982 | 0,0017 | 345 |
| Z70 | 18100 | 0,0026 | 147 |

| Tau peptide | sequence | $k_{on}$ (1/M.s) | $k_{off}$ (1/s) | Kd (nM) |
|---|---|---|---|---|
| Tau[273-318] | $_{273}$GKVQIINKKLDLSNVQSKCGSKDNIKHVPGGGSVQIV YKPVDLSKV$_{318}$ | 91914 | 0,0077 | 85 |

**Supplementary Figure 5 : Optimized mutant VHH Z70 has a better affinity for Tau than VHH E4-1.** Tables corresponding to $k_{on}$, $k_{off}$ and resulting Kd obtained from SPR experiments. Sequence of the Tau peptide used for the interaction with immobilized VHH Z70 on the chip is also included.

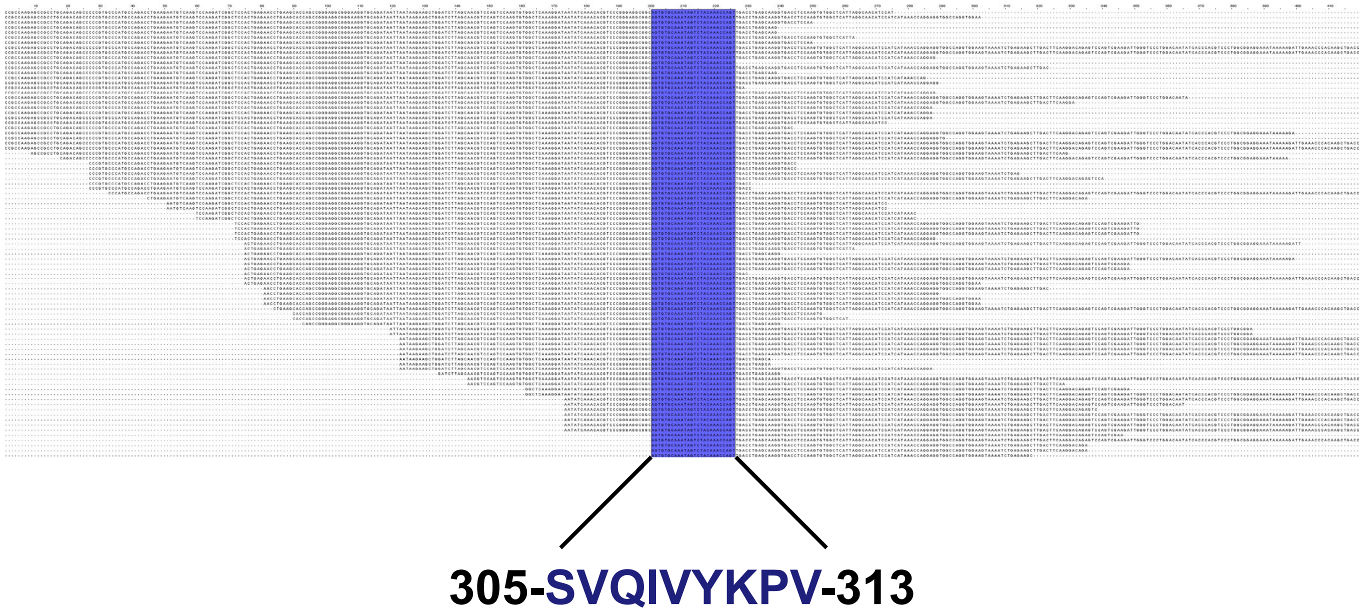



**Supplementary Figure 6 : Identification of the minimal epitope recognized by VHH Z70 using Tau fragment library and yeast two hybrid.** Sequence alignment of the 90 Tau fragments corresponding to the 90 positive colonies picked on selective growth conditions and thus binding VHH Z70. The minimal common sequence is highlighted. Sequences are not meant to be read.

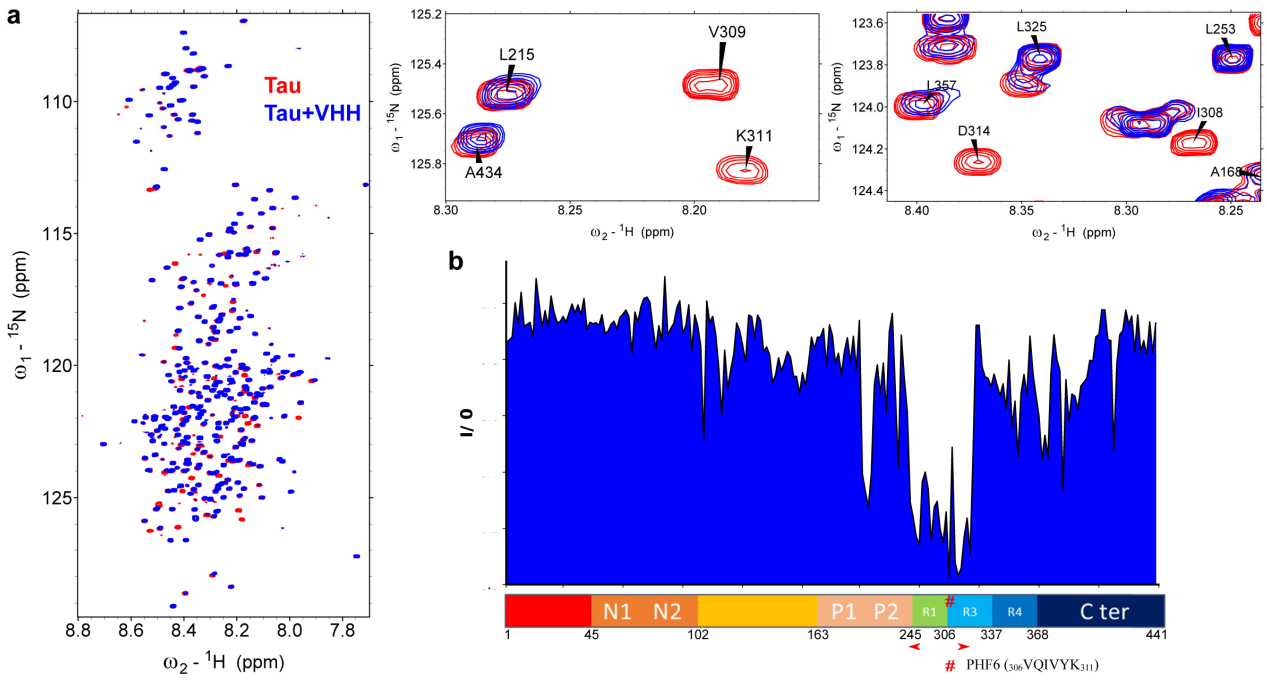


**Supplementary Figure 7 : Identification of VHH Z70 epitope using 2D HSQC NMR experiment. a** : Overlays of $^{1}$H, $^{15}$N, HSQC two-dimensional spectra enlargements of $^{15}$N-labelled Tau 2N3R, free (red) or mixed with non-labelled VHH Z70 (in blue). Spectra enlargements show broadened resonances corresponding to residues implicated in the interaction **b** : Intensities ratio I/I0 of corresponding resonances in the two-dimensional spectra of Tau 2N3R with equimolar quantity of VHH (I) or free in solution (I0) for residues along the Tau 2N3R sequence. Overlapping resonances are not considered (x-axis is not scaled). Tau 2N3R lacks the R2 repeats. Tau 2N3R residue numbering corresponds to the Tau 2N4R sequence, for clarity. The corresponding major broadened resonances, indicated by the red arrow heads, were mapped mostly on the PHF6 motif, showing PHF6 is sufficient for VHH Z70-Tau binding.

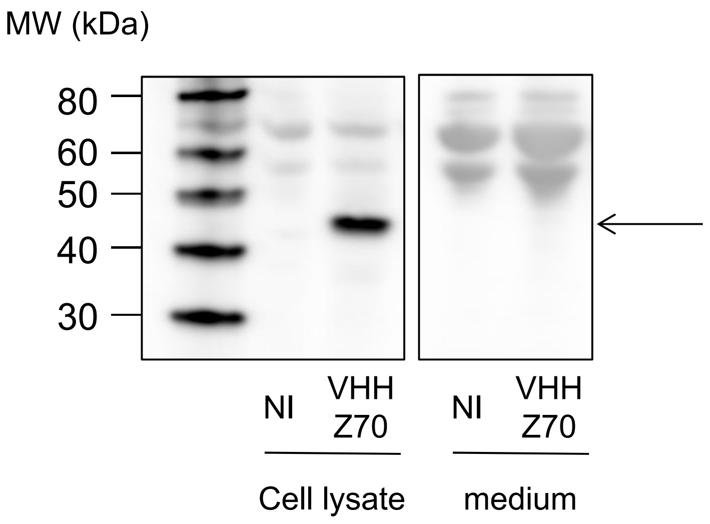


**Supplementary Figure 8 : Intracellular expression of VHH-Z70**: HEK293 cells were infected with LV encoding VHH-Z70 N-terminally fused to mCherry or non infected (NI). Forty-eight hours later, the cell lysate and the medium were recovered and analysed by western-blotting. VHH 70 expression (black arrow) was revealed thanks to the mCherry tag using primary antibody against mCherry.

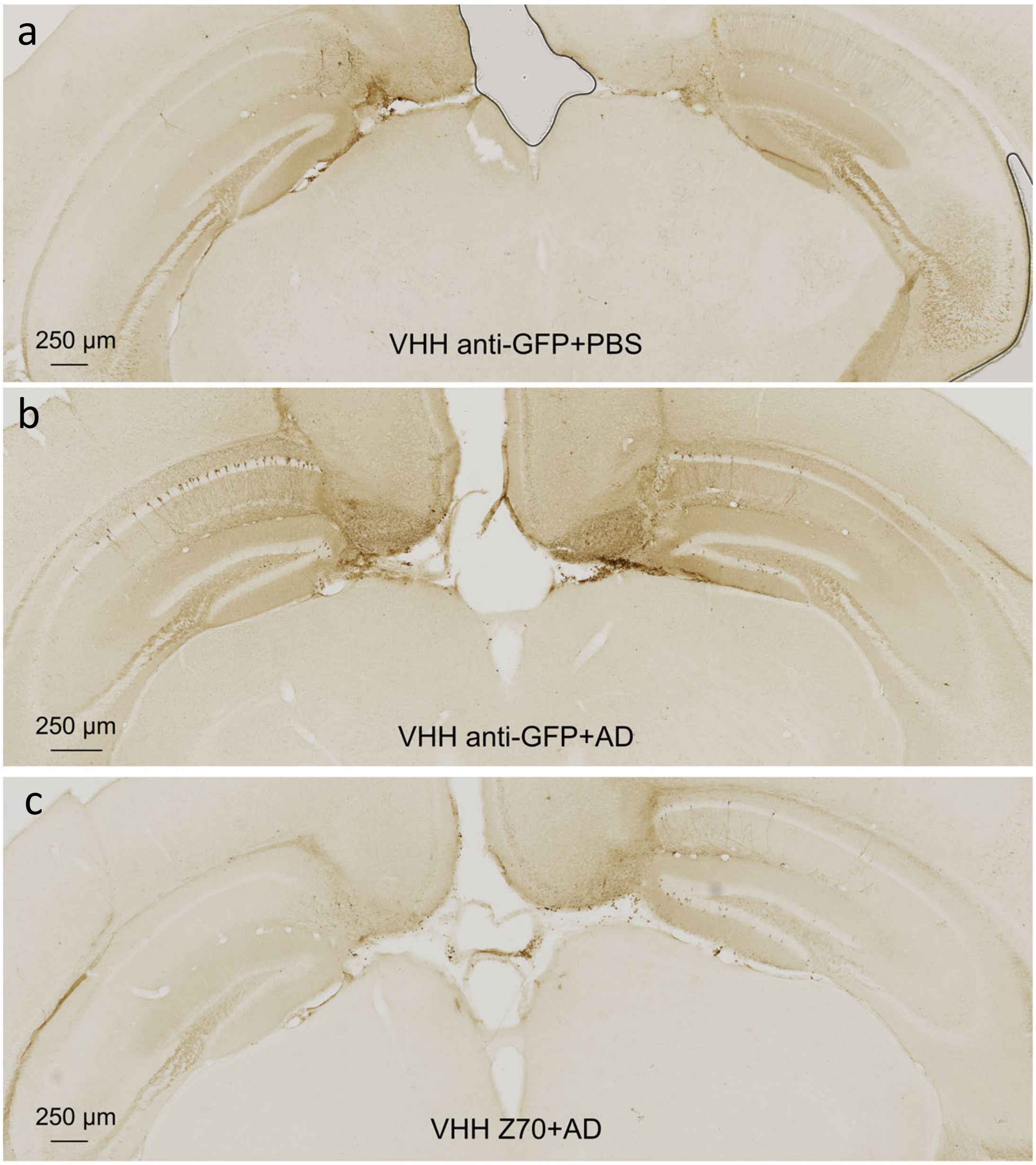


**Supplementary Figure 9: VHH Z70 reduces human Tau seeding induced by extracellular human pathological Tau species:** One month-old THY-tau30 mice were treated with bilateral injectionss of LVs encoding VHH-anti GFP (a,b) or VHH Z70 (c). Two weeks later mice received stereotaxic injection of PBS (a) or AD brain lysate (b,c). Mice were sacrificed 4 weeks later and the whole brains were processed for immunohistochemical analysis using AT8. Sections from the hippocampus (injection site) are shown. Scale bars are indicated on the figure.

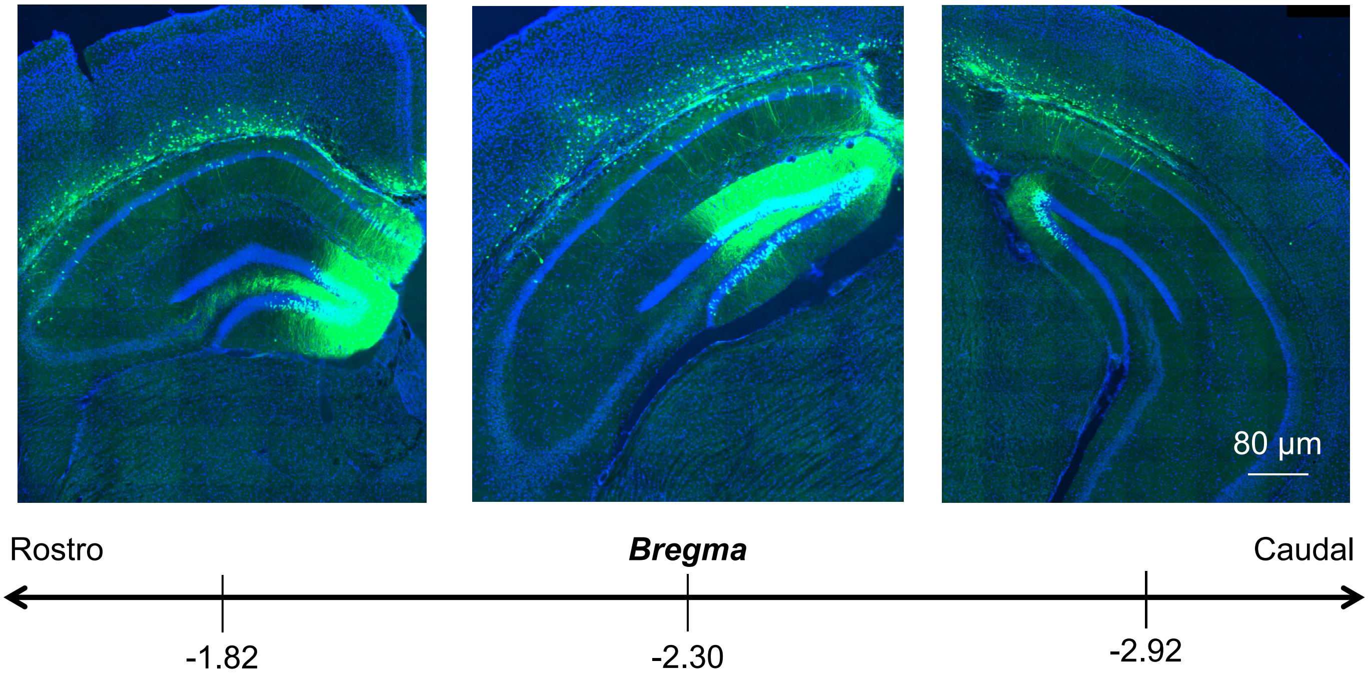


**Supplementary Figure 10: VHH Z70 expression in the hippocampus.** One month-old THY-tau30 mice were treated with bilateral intracranial injections of LVs encoding VHH Z70. Two weeks later mice received stereotaxic injection of AD brain lysate. Mice were sacrificed 4 weeks later and the whole brains were processed for immunohistochemical analyses. VHH Z70 was detected using a primary antibody against mCherry tag. VHH Z70-immunoreactivity is detected in all regions covering the bregma where tau pathology has been quantified.

3390
5pTCTGCACAATATTTCAAGCTATACCAAGCATACAATCAACTCCAAGCTAGAACCATGGCGGAAGTGCAGCTGCAGGCTC
3880
3pTCTTCTTTTTTGGAGGCTCGGGAATTAATTCCGCTTTATCCATCTTTGCGGCGGCCGCGCTACTCACAGTTACCTG

6690
5pCAGGGCAATAAAGTCGAACT,
6972
5pGACCTACAGGAAAGAGTTACTC

10829
5pCTATTCGATGATGAAGATACCCCACCAAACCCAAAAAAAGAGATCCTAGAACTAGACACTCTTTCCCTACACGACGCTCTTCC
10830
5pCCGGGCCTCTAGACACTAGCTACTCGAGGGGCCCCAGTGGCCCTATCTATGCGGCCGCTCAGACTGGAGTTCAGACGTGTGCTC

**Supplementary Fig. 11** Oligonucleotide sequences (Material and Methods)

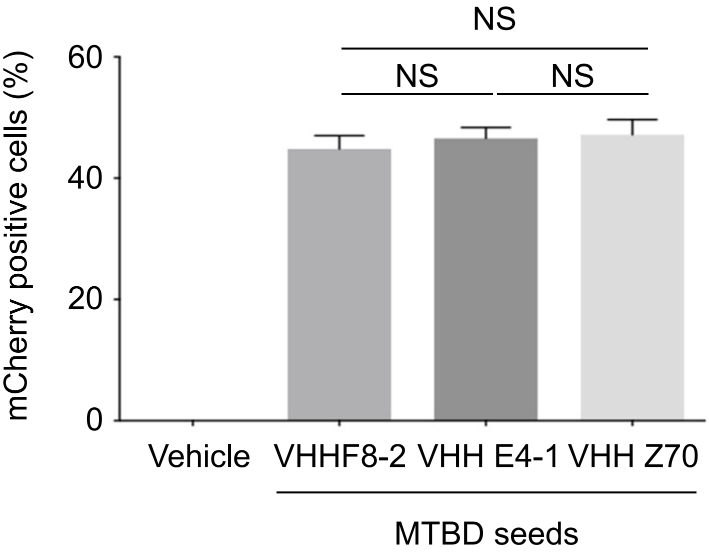


**Supplementary Fig. 12 : VHH expression in the biosensor seeding reporter cells.** HEK293 Tau RD P301S cells were transfected with plasmids encoding the different mCherry-VHH. mCherry fluorescence was evaluated by flow cytometry showing that transfection efficiency is equivalent : 44.8 (± 2.2%) for VHH-F8-2, 46.6 (± 1.8%) for VHH VE4-1 and 47.2 % (± 2;5%) for VHH Z70.
